# Computational investigation of IP_3_ diffusion

**DOI:** 10.1101/2022.08.19.504488

**Authors:** Roberto Ornelas Guevara, Diana Gil, Valérie Voorsluijs, Geneviève Dupont

## Abstract

Inositol 1,4,5-trisphosphate (IP_3_) plays a key role in calcium signaling. After stimulation, it diffuses from the plasma membrane where it is produced to the endoplasmic reticulum where its receptors are localized. Based on *in vitro* measurements, IP_3_ was long thought to be a global messenger characterized by a diffusion coefficient of ~280 µm^2^s^−1^. However, *in vivo* observations revealed that this value does not match with the timing of localized Ca^2+^ increases induced by the confined release of a non-metabolizable IP_3_ analog. A theoretical analysis of these data concluded that in intact cells diffusion of IP_3_ is strongly hindered, leading to a 30- fold reduction of the diffusion coefficient. Here, we performed a new computational analysis of the same observations using a stochastic model of Ca^2+^ puffs. Our simulations concluded that the value of the effective IP_3_ diffusion coefficient is close to 100 µm^2^s^−1^. Such moderate reduction with respect to *in vitro* estimations quantitatively agrees with a buffering effect by non-fully bound inactive IP_3_ receptors. The model also reveals that IP_3_ diffusion is not much affected by the endoplasmic reticulum, which represents an obstacle to the free displacement of molecules, but can be significantly increased in cells displaying elongated, 1-dimensional like geometries.

## Introduction

Within all cell types, Ca^2+^ signaling is controlled by a variety of channels and pumps that allow for rapid and highly regulated Ca^2+^ fluxes at specific locations of the cell (Bootman et al., 2001). In many instances, Ca^2+^ increases are initiated by the formation of inositol 1,4,5-trisphosphate (IP_3_) at the plasma membrane (Berridge, 1997). IP_3_ diffuses in the cytoplasm and binds its receptors located on the endoplasmic reticulum that contains a large quantity of rapidly mobilizable Ca^2+^ (Sammels et al., 2010). Activity of these receptors is also regulated by cytosolic Ca^2+^, both positively and negatively (Bezprozvanny et al., 1991). At the spatial level, IP_3_ receptors (IP_3_R) are not homogeneously scattered on the ER membrane, but rather grouped in clusters of ~10-20 channels (Swillens et al., 1999; Smith and Parker, 2009a). Because the number of clusters in cells is limited, coupling between clusters has been much investigated to understand how Ca^2+^ signaling can be coordinated at the cellular level, giving rise to Ca^2+^ oscillations and waves (Bootman et al., 1997; Skupin et al., 2008; Voorsluijs et al., 2019). Most studies have focused on Ca^2+^-mediated communication between clusters, since it was assumed that all clusters rapidly experience the increase of IP_3_ that results from the stimulation of the cell. In cytosolic extracts of *Xenopus* oocytes, apparent diffusion coefficients of Ca^2+^ and IP_3_ indeed equal 38 ± 11 µm^2^s^−1^ and 283 ± 53 µm^2^s^−1^, respectively (Allbritton et al., 1992). Considering in addition the respective rates of Ca^2+^ removal from the cytoplasm and IP_3_ metabolism, these values led Allbritton et al. (1992) to conclude that Ca^2+^ mostly acts in restricted domains and that IP_3_ is a global messenger.

The notion that IP_3_ acts as a global messenger was however contradicted by indirect *in vivo* observations (Dickinson et al., 2016). To evaluate the rate at which IP_3_ diffuses in an intact cell, the group of Ian Parker used the IP_3_-evoked liberation of Ca^2+^ from a cluster of IP_3_Rs as a detector of the presence of IP_3_ at the cluster location. Such Ca^2+^ increases evoked by clusters of IP_3_Rs are well-known as Ca^2+^ puffs (Yao et al., 1995). If a non-metabolizable IP_3_ analog is released at one extremity of an elongated SH-SY5Y cell, the time laps between the localized IP_3_ increase and the occurrence of the first Ca^2+^ puff significantly increases with the distance between the spot of IP_3_ release and the location of Ca^2+^ rise. This time laps, called *latency*, reflects the time taken by IP_3_ to diffuse on this distance. Accordingly, when the IP_3_ analog is uniformly released across the entire cell, latency does not show any systematic variation along the cell length but decreases with increasing IP_3_ concentration. The fact that latency increases with the distance from the IP_3_ release spot clearly indicates that IP_3_ does not act as a global messenger in these conditions. From these observations, Dickinson et al. (2016) inferred the value of the IP_3_ diffusion coefficient by resorting to a simplified mathematical expression for the probability of Ca^2+^ puff occurrence coupled to 1-dimensional (1D) simulations of IP_3_ diffusion. These calculations predicted an effective IP_3_ diffusion coefficient lower than 10 µm^2^s^−1^, *i*.*e*. about 30 times slower than *in vitro* estimations.

This slowing down of IP_3_ diffusion within cells as compared to cytosolic extracts was ascribed to the presence of IP_3_Rs that are not fully bound to IP_3_ (Dickinson et al., 2016; Taylor and Konieczny, 2016). IP_3_Rs are indeed tetramers that release Ca^2+^ only when each IP_3_R monomer is occupied by IP_3_ (Alzayady et al., 2016). Given that IP_3_ binding on IP_3_Rs is not cooperative, most of the receptors are partially bound as long as [IP_3_] remains lower than the K_D_ of IP_3_ binding, *i*.*e*. ~100 nM (K_D_=119 nM reported by Taylor and Konieczny, 2016) and thus act as a buffer of IP_3_. Assuming fast binding and unbinding of IP_3_ to and from its receptor, the resulting effective diffusion coefficient (Dupont et al., 2016) is given by:

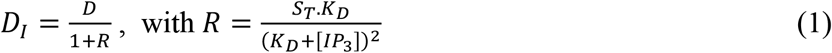

where *D* represents the IP_3_ diffusion coefficient in the absence of buffers, *S*_*T*_ the concentration of IP_3_R monomers, and *K*_*D*_ the equilibrium dissociation constant of IP_3_ from its receptor. *S*_*T*_ is cell dependent, in the range of 80 nM to 2 µM (Spät et al., 1986; Wojcikiewicz, 1995; Dupont et al., 2008; Taylor and Konieczny, 2016) with an estimated value of 542 nM in SH-SY5Y cells (Wojcikiewicz, 1995). Thus, the buffering effect described by equation (1) is expected to induce a reduction of the effective diffusion coefficient of ~2.25 at [IP_3_] = *K*_*D*_. In Dickinson et al. (2016) experiments, [IP_3_] was in the Ca^2+^ oscillatory range. Because IP_3_ concentrations are typically of the order or lower than the K_D_ of the IP_3_R in the oscillatory range (Tanimura et al., 2009), one can estimate that the effective diffusion coefficient of IP_3_ considering IP_3_ buffering by not fully bound receptors (*D*_*I*_) should be *~*100 µm^2^s^−1^ (Figure 1). Such discrepancy led Dickinson et al. (2016) to hypothesize the existence of two distinct populations of IP_3_Rs, with the most active exhibiting different binding and/or gating properties. These different populations were proposed to be the molecular bases of the observed two modes of Ca^2+^ release: one early pulsatile Ca^2+^ activity at puff sites and one later, spatially more diffuse Ca^2+^ liberation (Lock and Parker, 2020).

**Figure 1.**
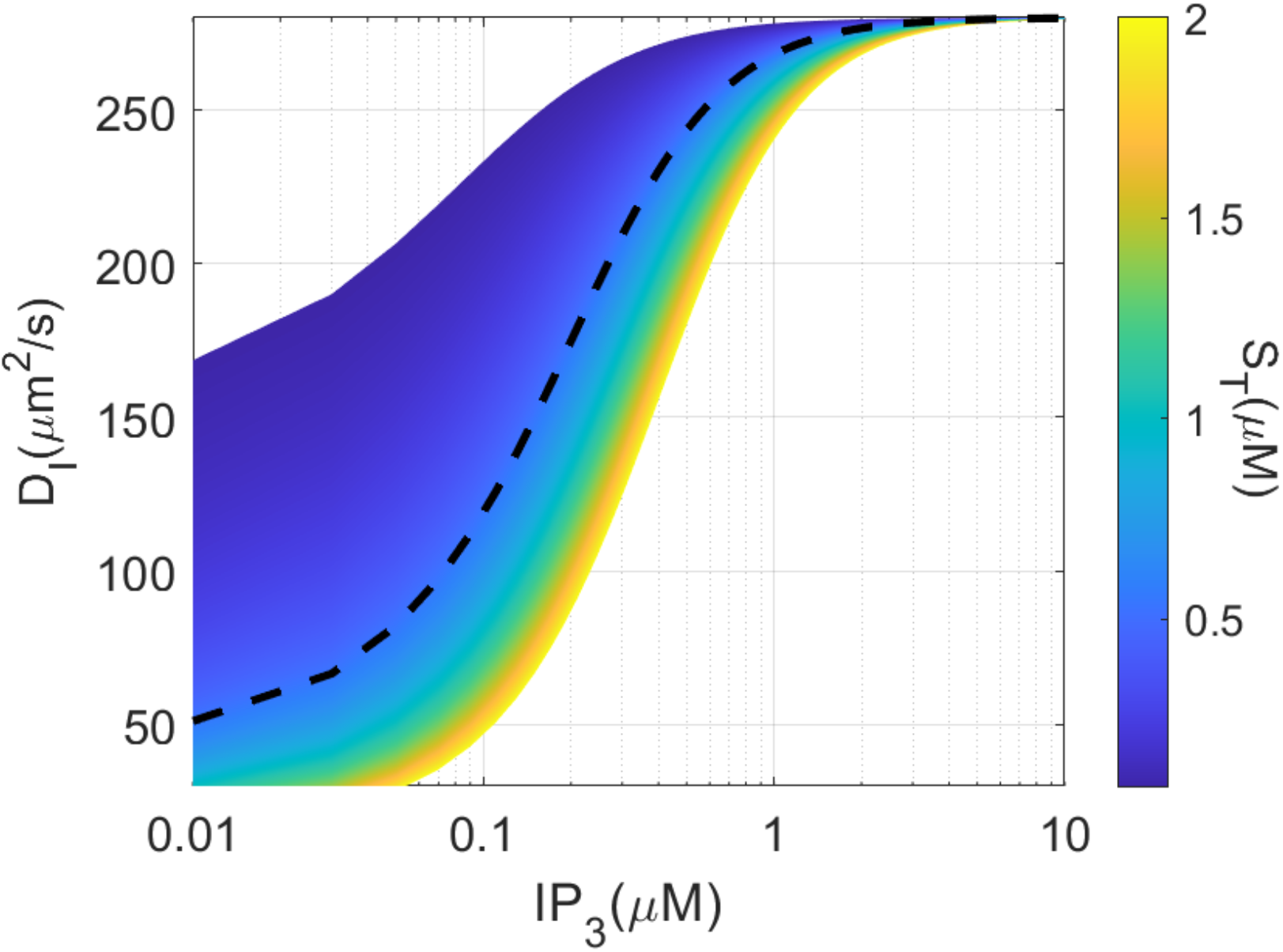
Theoretical values of the effective IP_3_ diffusion coefficient depending on the concentration of IP_3_ and of the monomers of IP_3_ receptors (S_t_). The dashed line shows this diffusion coefficient as a function of [IP_3_] for S_t_ = 542 nM, which is the value estimated for SH-S5Y5 neuroblastoma cells. Values have been calculated using Equation (1). See text for details.

Besides, a value of *D*_*I*_ lower than 10 µm^2^s^−1^ raises questions about some observations related to Ca^2+^ waves. For example, in ascidian eggs that have a radius between 25 and 38 µm, Ca^2+^ waves propagate across the entire fertilized egg in less than 10 s while IP_3_ is only synthesized at the plasma membrane (Dupont and Dumollard, 2004). The question is even more critical for intercellular Ca^2+^ wave, which in many cases rely on the propagation of IP_3_ through gap junctions (Decrock et al., 2018). Intracellular diffusion of IP_3_ with a coefficient ≤ 10 µm^2^s^−1^ could not account for waves propagating at rates ≥ 10 µms^−1^ among a cell population (Dupont et al., 2007; Leybaert, 2016).

The aim of the present study is to re-investigate the characteristics of IP_3_ diffusion using a stochastic model that explicitly simulates Ca^2+^ puff dynamics and allows a realistic computational description of the experiments performed by Dickinson et al. (2016). Based on the observations of puff latencies performed by these authors, we propose a new computational treatment to infer the value of the effective diffusion coefficient of IP_3_, leading to different conclusions. The reasons for the different outcomes between the two studies are analyzed in the discussion. We also investigate how cell geometry and the presence of the ER membranes, which provide an obstacle to the free displacement of molecules, affect the rates at which IP_3_ diffuses in a cell-like environment.

## Results

### Methodology

We performed explicit stochastic simulations of Ca^2+^ releasing activities of puff sites located at different distances from an IP_3_ source, in order to directly simulate the experimental protocol that was used to estimate the IP_3_ diffusion coefficient *in vivo* (Dickinson et al., 2016). The mathematical model is based on a previously proposed fully stochastic description of the Ca^2+^ exchanges between the ER and the cytosol via IP_3_Rs, SERCA pumps and a leak from the ER (Voorsluijs et al., 2019). In this work, SERCA pumps, Ca^2+^ leakage and Ca^2+^ diffusion are described deterministically. The effective value of Ca^2+^ diffusion coefficient taking Ca^2+^ buffering into account is considered, *i*.*e*. 40 µm^2^s^−1^ (Allbritton et al., 1992). To simulate Ca^2+^ puffs, the release of Ca^2+^ via the IP_3_Rs is described stochastically, using the Gillespie’s algorithm. Each cluster of IP_3_Rs (Figure S1) is described as a whole and can be in four states: one open (O), one closed (C) and two different inhibited ones (I_1_ and I_2_). These states describe the global behavior of a cluster composed of close-by IP_3_Rs. The phenomenological model was shown to reproduce experimentally observed statistical properties of Ca^2+^ puffs (Calabrese et al., 2010) and to describe the passage from localized puffs to global Ca^2+^ spikes when clusters are effectively coupled by Ca^2+^ diffusion (Voorsluijs et al., 2019).

To evaluate the effective diffusion coefficient of IP_3_, this model is extended to take the dependence of puff occurrence on IP_3_ concentration into account. Thus, in the Gillespie’s simulations, the propensity of transition of the cluster from the closed (C) to the open (O) state now writes:

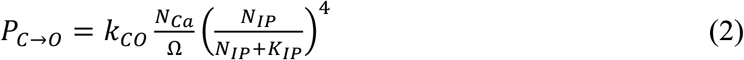

where *k*_*CO*_ stands for the rate constant that characterizes the passage of the cluster from the closed to the open state, *N*_*Ca*_ is the number of cytosolic Ca^2+^ ions and Ω is the extensivity parameter. *N*_*IP*_ and *K*_*IP*_ represent the number of IP_3_ molecules and the IP_3_ dissociation constant of the IP_3_R (multiplied by Ω), respectively. Although the model does not simulate individual IP_3_ receptors, we assumed that puff firing probability is related to the probability of one tetrameric IP_3_R to be fully bound to IP_3_ (Taylor and Konieczny, 2016). As shown below, Equation 1 indeed allows to reproduce the exponential dependence of mean first puff latency on IP_3_ concentration reported experimentally (Dickinson et al., 2012; 2016).

The evolution of IP_3_ concentration is described deterministically and two different protocols of photorelease of caged IP_3_ are simulated. The first one corresponds to a spot photorelease of caged IP_3_ in a small region of the cell:

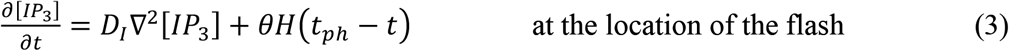

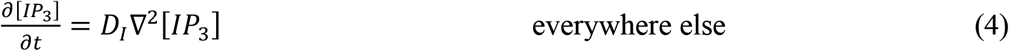

In the second one, called “distributed photorelease”, IP_3_ is liberated at different spots to get an increase in [IP_3_] that is nearly spatially homogeneous in the whole cell. In this case:

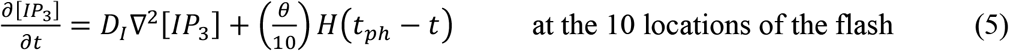

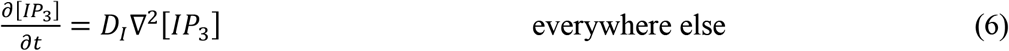

In Equations 3-6, *D*_*I*_ stands for the effective IP_3_diffusion coefficient, θfor the rate of IP_3_ release upon laser flash and *H* for Heaviside function. This function equals 1 if t ≤ t_ph_ and 0 otherwise. Considering that θ is determined by the intensity of the flash in the spot photorelease case (Equation 3), a ten times smaller value is used to simulate distributed photorelease (Equation 5) because the same total amount of IP_3_ is distributed among the ten IP_3_ releasing sites. Basal IP_3_ concentration is set to 50 nM (Luzzi et al., 1998).

Except for the process describing IP_3_ dynamics listed here above, the values of parameters are taken from Voorsluijs et al. (2019) and are listed in Table Supplement 1.

### Validation of the model

Low concentrations of IP_3_ typically evoke local Ca^2+^ signals known as Ca^2+^ puffs (Yao et al., 1995; Tovey et al., 2001). As the concentration of IP_3_ increases, Ca^2+^ puffs become more frequent and transform into Ca^2+^ waves spreading regeneratively across the cell. These repetitive waves, also known as Ca^2+^ spikes, are more regular than puffs and are often referred to as Ca^2+^ oscillations. Their stochastic origin is visible by the linear relation between the variance on the interspike interval and the mean interspike interval itself (Skupin et al., 2008; Thurley et al., 2014). In experiments, Ca^2+^ waves propagation can be hindered by loading the cells with the slow Ca^2+^ buffer EGTA, which reduces communication between puff sites (Dargan and Parker, 2003). At high EGTA concentrations, each cluster practically behaves as an independent entity and thus generates Ca^2+^ puffs on a large range of [IP_3_].

Such behavior is well reproduced by the stochastic model, as seen by simulations in a simplified 2D geometry (Figure 2A-D). Four different conditions were used: low and intermediate IP_3_ concentration, with and without coupling between clusters by Ca^2+^ diffusion. In all cases, [IP_3_] is constant in time and space. To model puff dynamics when clusters are uncoupled (corresponding to the presence of EGTA), a single cluster was simulated and the local Ca^2+^ concentration averaged in 1fL around the cluster was monitored. The resulting dynamics of [Ca^2+^] (Figure 2 A-B) corresponds to Ca^2+^ puffs, with statistical properties in good agreement with observations and an average inter-puffs interval that decreases with [IP_3_] (Voorsluijs et al., 2019). The mean interpuff intervals are 4.60 ± 4.25 s and 1.86 ± 1.79 at 0.15 µM and 0.38 µM [IP_3_], respectively. To model spikes dynamics when clusters are coupled by diffusion, 10 clusters of IP_3_R were randomly distributed in a 5 µm x 5 µm system and the global [Ca^2+^] averaged over the whole system was monitored (Figure 2C-D). The mean interspike interval and the coefficient of variation (CV) decrease when increasing [IP_3_], as reported previously (Thurley et al., 2014): the mean interspike intervals are 129.6 ± 64.00 s and 108.26 ± 49.22 s at 0.15 µM and 0.38 µM [IP_3_], respectively. To simulate the spatio-temporal profiles of puffs and spikes, simulations were performed in a 50 µm x 10 µm system, while keeping the same average density of clusters (Figure 2E and 2F). While a low, homogeneous concentration of IP_3_ induces random puff activity, a large bolus of IP_3_ released at one extremity of the cell initiates a global Ca^2+^ spike that propagates as a Ca^2+^ wave.

**Figure 2.**
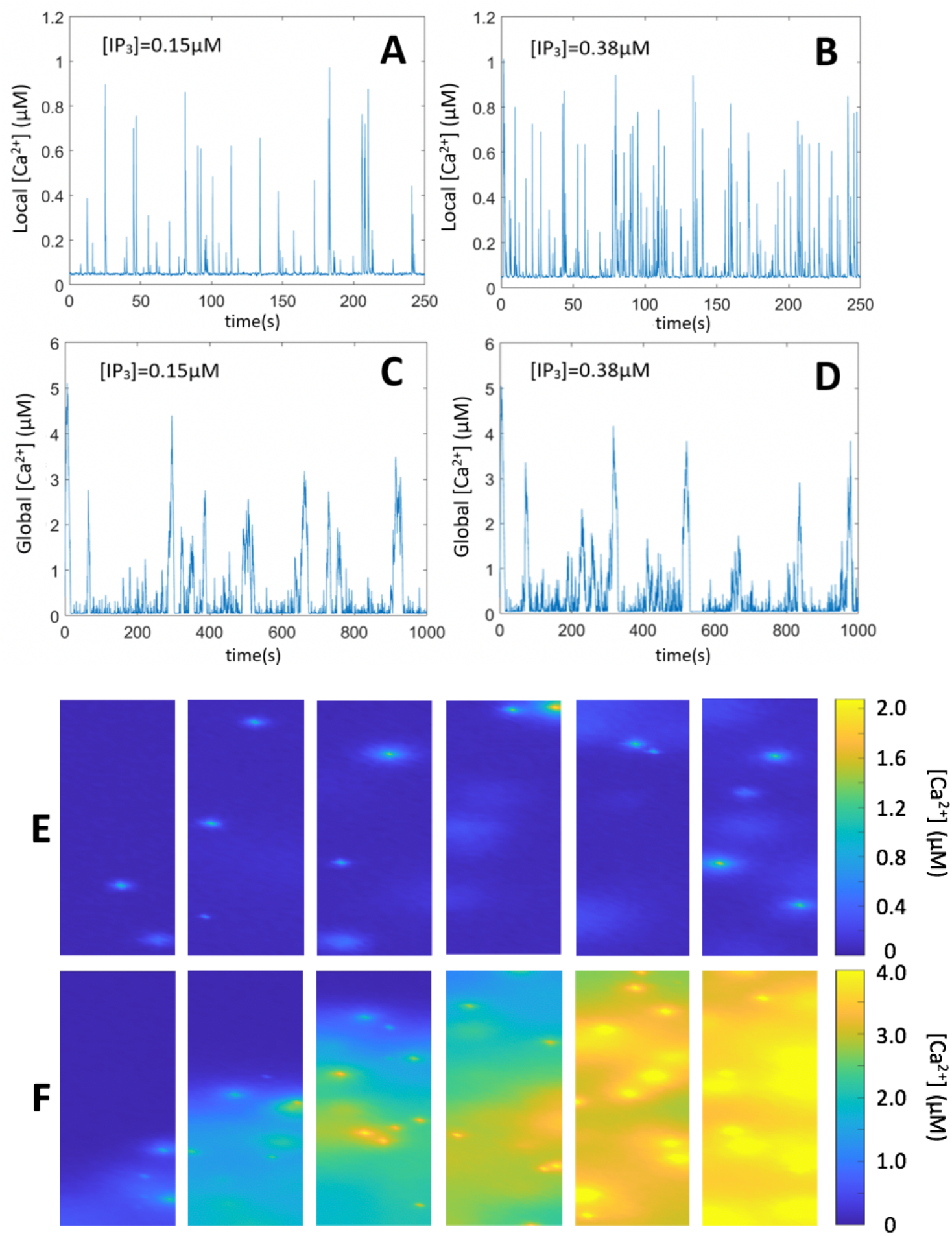
Stochastic simulations of Ca^2+^ puffs and spikes. **A and B** show the evolution of Ca^2+^ concentration in a 1 fL volume centered around the cluster at two [IP_3_]. For the 2 panels, one cluster, located at the centre of a 5 × 5 µm^2^ 2D system is considered. For panels **B and C**, a larger 5 × 10 µm^2^ system containing 10 clusters is simulated. All values of parameters are the same as in A and B and listed in Table S1. Time series show the evolution of Ca^2+^ concentration averaged on the whole system. Panels **E and F** show the spatio-temporal evolutions of Ca^2+^ puffs and spikes, respectively. In the two cases, the system is 50 × 10 µm^2^ large and 10% of the total surface is occupied by clusters (200 clusters). For panel E, [IP_3_] is constant and equal to 0.075 µM everywhere in the cell. For panel F, IP_3_ was released in one spot (with dimensions 0.5 × 0.5 µm^2^) located in the bottom center of the simulated cell. For panels E and F, times corresponding to each subpanels are: 0.1, 0.6, 2.5, 3.4 and 5.2 s after the increase in [IP_3_] from 10 nM. In panels A-E, [IP_3_] is increased at time 0, and equations (3) and (5) are not considered. In panel F, the release of IP3 is simulated using equation (3), with θ= 1000µMs^.1^ and t_ph_ = 500 ms.

Upon a spatially uniform increase in IP_3_ concentration, latencies of first puff occurrence are exponentially distributed (Yao et al., 1995; Dickinson et al., 2012). Such distributions, obtained by model simulations, are shown in Figure 3A-C, together with their means and associated standard error of the mean (SEM) values. These results were used to infer the values of the rate of release of caged IP_3_ in response to the flash, *i*.*e*. the value of parameter *θ* in Equations 3 and 5. Starting from the relation between mean first puff latency and flash duration reported by Dickinson et al. (2016) for the distributed photorelease of IP_3_ - and replotted in Figure 3D-, we seek the values of IP_3_ concentrations that, in the simulations, gave the same mean latencies as those reported experimentally using Figure 3C. Next, we numerically evaluated the value of *θ* allowing to reach these spatially uniform concentrations of IP_3_ when assuming 10 photorelease spots of durations equal to 0.1, 0.2 and 0.5 s respectively, as in the experiments. Good agreement between mean latencies and flash durations was found for *θ*=600 µM/s (Figure 3D).

**Figure 3.**
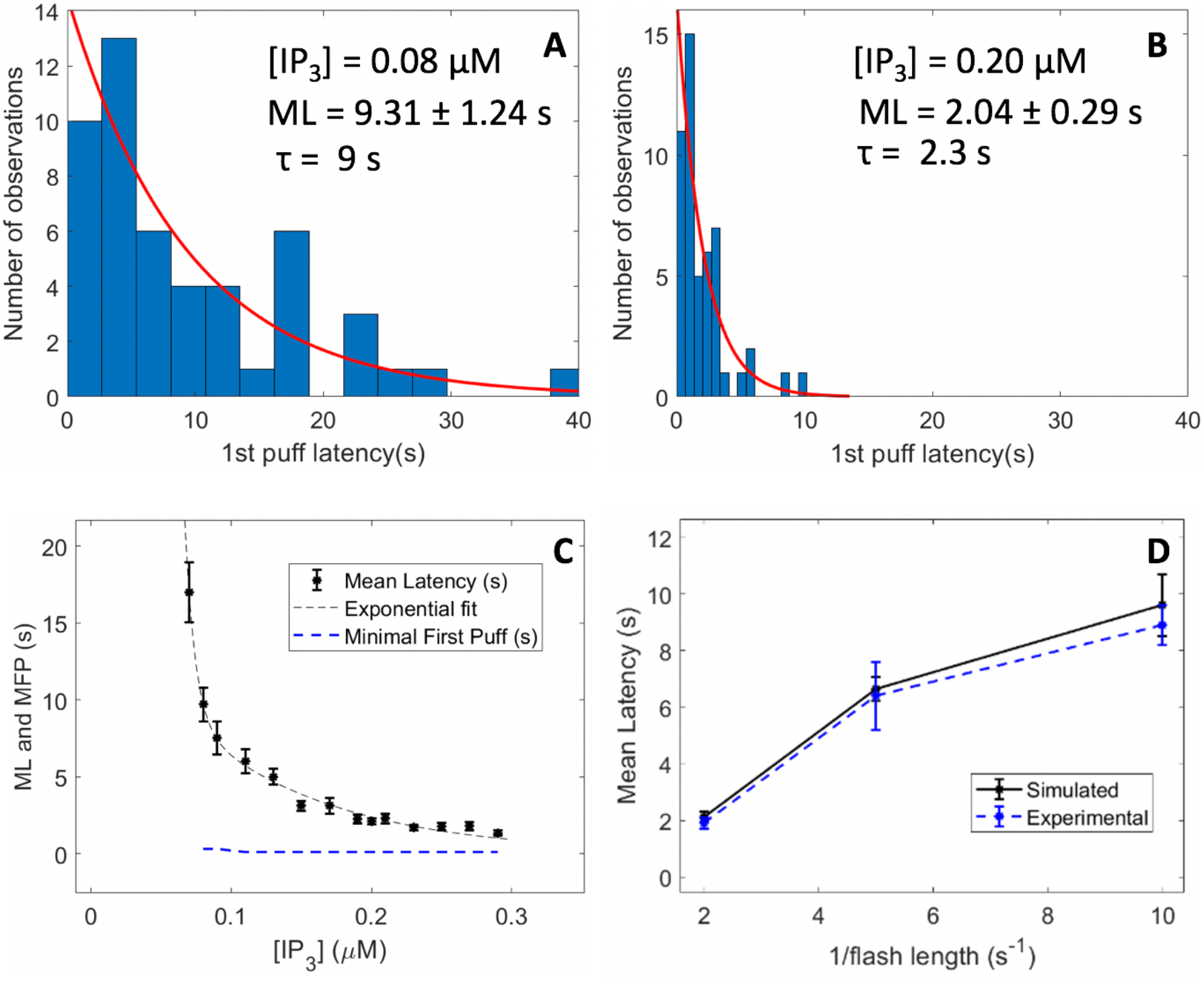
Statistics of the latencies of first puffs simulated with the model. Panels **A and B** show histograms of first puff latencies resulting from simulations of one cluster site in a square 5 × 5µm^2^ 2D geometry. A puff is defined as an increase in the cytosolic Ca^2+^ concentration in a 1 fL volume centered around the cluster that exceeds 0.1 µM. First puff latencies show an exponential distribution with characteristic decrease times matching experimental observations. Time t = 0 corresponds to the moment of [IP_3_] increase from 50nM to indicated values. The black stars in panel **C** show the mean first puff latencies (ML) as a function of [IP_3_]. The blue dashed line shows the minimal first puffs (MFP), *i*.*e*. the time at which the first puff occurred. Minimal first puffs are practically independent of [IP_3_] and are always close to 300 ms. For each [IP_3_], 50 independent simulations were run. In panels A, B and C, [IP_3_] is increased at time 0, and equations (3) and (5) are not considered. In panel D, photorelease of caged IP_3_ is also simulated, using equation (5). Best fit with the observations of Dickinson et al. (2016) was found when considering *θ* = 600 µMs^−1^. This value was found by looking for the value of *θ* that allows to obtain the steady states [IP_3_] leading to the ML’s corresponding to the flash durations (t_ph_) used in Dickinson et al.’s experiments, i.e. 8.9 ± 0.5 s, 6.4 ± 1.2 s and 1.9 ± 0.2 s for the 0.1, 0.2 and 0.5 s flash durations, respectively. Error bars indicate ± SEM.

When caged IP_3_ is photoreleased at one extremity of the cell, puffs begin on average after longer latencies at greater distance from the spot (Dickinson et al., 2016). This reflects the time taken for IP_3_ to diffuse up to the cluster, and IP_3_ dilution. Dickinson et al. (2016) used mean first puff latencies at different distances from the photorelease spot to estimate the effective diffusion coefficient of IP_3_. In addition to *mean* first puff latencies, the time interval between the flash and the observation of the first puff at a given distance from the release site is also a relevant quantity. We called this latency “*minimal* first puff latency”. It can be regarded as the time laps during which all clusters located at a given distance from the flash site remain silent, indicating that IP_3_ diffusion up to this point has been negligible. As visible in the distribution of puff latencies upon a global IP_3_ increase, the minimal first puff latency is shorter than 0.5 s as soon as [IP_3_] exceeds 80 nM (Figure 3C, blue dashed line). In agreement with this, the minimal first puff latency is not affected by the duration of the flash in the case of distributed photorelease of IP_3_ (Dickinson et al., 2016). If the source of IP_3_ is spatially restricted, minimal first puff latency provides a reliable indication of the time at which IP_3_ has started to increase at a given location since IP_3_ diffusion can be viewed as a deterministic process and since the number of puff sites analyzed is large enough. Moreover, as discussed below, the minimal first puff latency is not influenced by the possible uncomplete Ca^2+^ buffering by EGTA, which could accelerate puff triggering. Indeed, first puffs arise in conditions where most, if not all, nearby clusters are inactive.

### Effective diffusion coefficient of IP_3_

We investigated the relation between first puff latency and distance from spot photorelease by performing independent simulations for twelve clusters, each one located at a different distance from the photorelease spot in an ellipse-shaped 2D cell. Such a configuration allows to reproduce the absence of communication between the cluster sites that results from the addition of EGTA. The ellipse shape was chosen instead of the rectangle used in the previous figures to avoid artefactual effects in the corners. Each simulation was run up to the opening of the cluster and this time was then monitored. The same simulation was performed 50 times for each distance between the spot and the cluster. As shown in Figure 4 (upper row), minimal first puffs latencies (black stars) simulated with an effective IP_3_ diffusion coefficient of 100 µm^2^s^−1^ –which corresponds to the expected value given the IP_3_ buffering capacity of the cytosol of SH-SY5Y cells, as discussed in the Introduction– are in agreement with experimental observations of Dickinson et al. (2016). Thus, the time taken by IP_3_ to diffuse from the photorelease spot to the locations of the clusters is such that the relation between the distance from the flash spot and the duration of the period of inactivity are well reproduced. Agreement between simulated and observed mean first puff latencies is limited to the case of the 0.1 s flash, *i*.*e*. the lowest IP_3_ concentration. For larger IP_3_ concentrations (0.2 and 0.5 s flash), we reasoned that the discrepancy between the idealized simulations of cells containing one single cluster and experiments may be due to a slight stimulation of puff activity by Ca^2+^ in the experiments, which is not totally buffered by EGTA. Because we only model one cluster at a time, this effect does not occur in our simulations.

**Figure 4.**
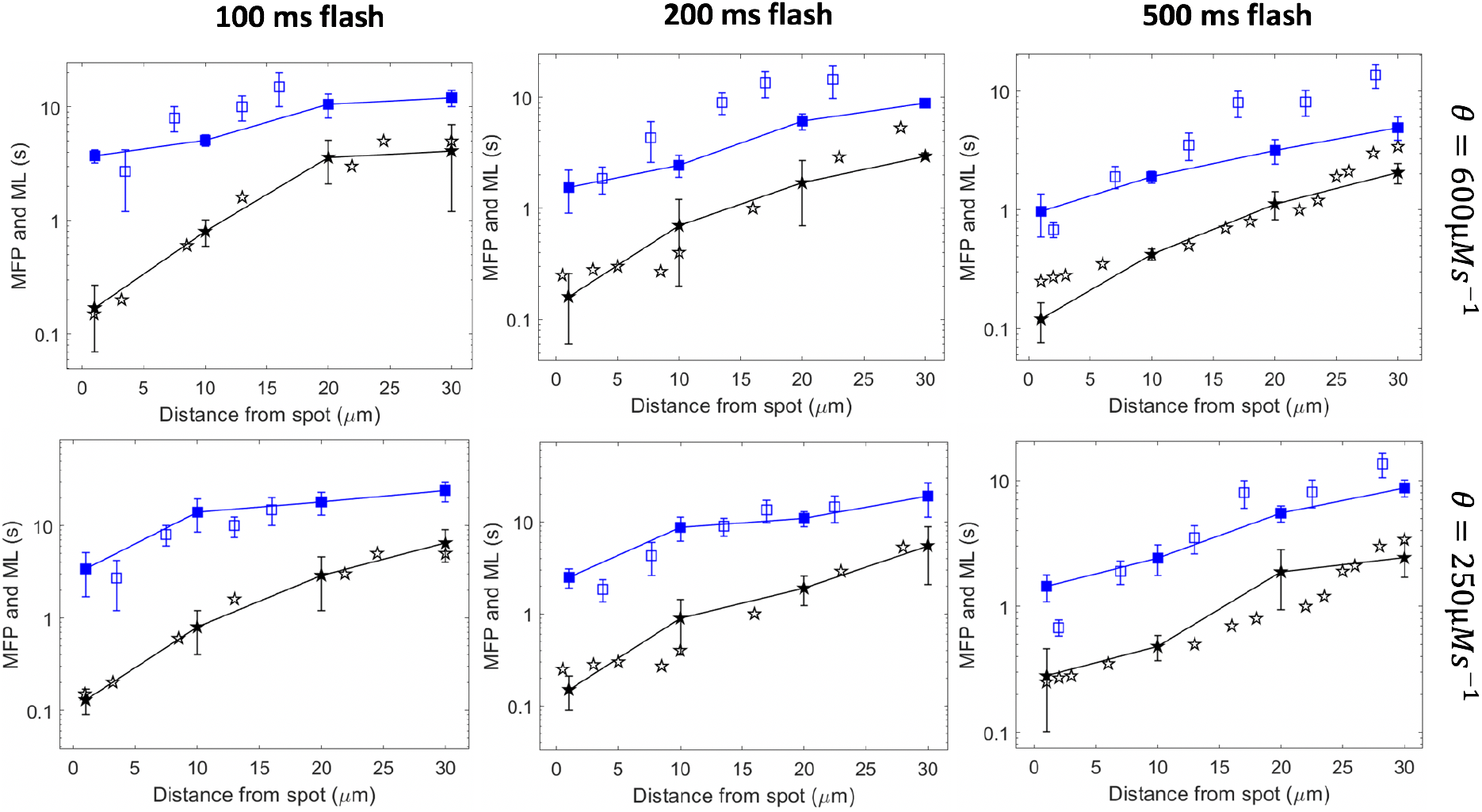
Mean latencies (ML) and minimal first puff latencies (MFP) as simulated with an effective IP_3_ diffusion coefficient equal to 100 µm^2^s^−1^ and a rate of IP_3_ increase at the photorelease spot of 600 µMs^−1^ (upper row) or 250 µMs^−1^ (bottom row). In all panels, square represent mean latencies and stars, minimal first puff latencies. Plain symbols are theoretical predictions while empty symbols are the experimental observations of Dickinson et al. (2016). Lines are drawn between simulation results. Simulations are performed in a 2D ellipse-shaped geometry with the spot of IP_3_ release occurring in a 0.25 µm radius circle at one extremity of the simulated cell. For each theoretical point, 50 independent simulations were run, considering one cluster at a time. Error bars indicate ± SEM. For minimal first puffs, the 50 simulations were divided in 10 groups of 5 simulations among which the minimal first puff was considered.

In line with this assumption, a lower value of parameter *θ* (250 µMs^−1^) representing the rate of IP_3_ release due to the flash (Equations 3 and 5) allows to get a good agreement both for the mean and the minimal first puff latencies (Figure 4, lower row). Best agreement is obtained with *D*_*I*_ = 100 µm^2^s^−1^, as compared to smaller or larger values (Supplemental Figure S2A-B). This value of *θ* also allows us to reproduce the experimentally observed dependency of the mean latency on the flash duration when considering a higher basal Ca^2+^ level in the simulations (Supplemental Figure S2C). Our results thus suggest that the value of the rate of IP_3_ photorelease (parameter *θ*) directly inferred from the experiments of distributed photorelease is overestimated because the slight increase in cytosolic Ca^2+^ occurring at 5 µM EGTA accelerates puff occurrence. Considering this, simulations are in good agreement with experimental observations concerning mean and minimal first puff latencies with the value of the IP_3_ diffusion coefficient taking IP_3_ binding to the receptors into account, *i*.*e*. 100 µm^2^s^−1^. The latter value also allowed to reproduce the relation between puff latencies and distance from the flash measured in COS-7 cells (Supplemental Figure S3).

### Influence of cell shape

Up to this point, simulations have been performed in 2D, rectangular or ellipse-shaped systems. However, the 3D character and the specific geometry of the cells are expected to influence the rate at which IP_3_ propagates into the cytoplasm through diffusion. To address this question using computational simulations (Figure 5), we first considered an ellipsoid (50 × 10 × 5 µm) in which 20 cluster sites are located randomly at a distance shorter than 0.1 µm from the plasma membrane (Smith et al., 2009). IP_3_ is assumed to be released in a 0.25 µm radius sphere located at the left extremity of the cell. The rate of IP_3_ increase (parameter *θ* in Equations 3 and 5) was adapted according to the change in the cytoplasmic volume. Two seconds after the simulated flash, a gradient of IP_3_ is established (Figure 5). Consequently, the dynamics of puff activity is rather different depending on the distance from the flash (compare the Ca^2+^ time series in Figure 5). Agreement between minimal and mean first puff latencies computed in simulations with an effective diffusion coefficient of IP_3_ equal to 100 µm^2^s^−1^ and observations of Dickinson et al. (2016) is slightly improved in this 3D configuration as compared to the 2D situation (Supplemental Figure S4).

**Figure 5.**
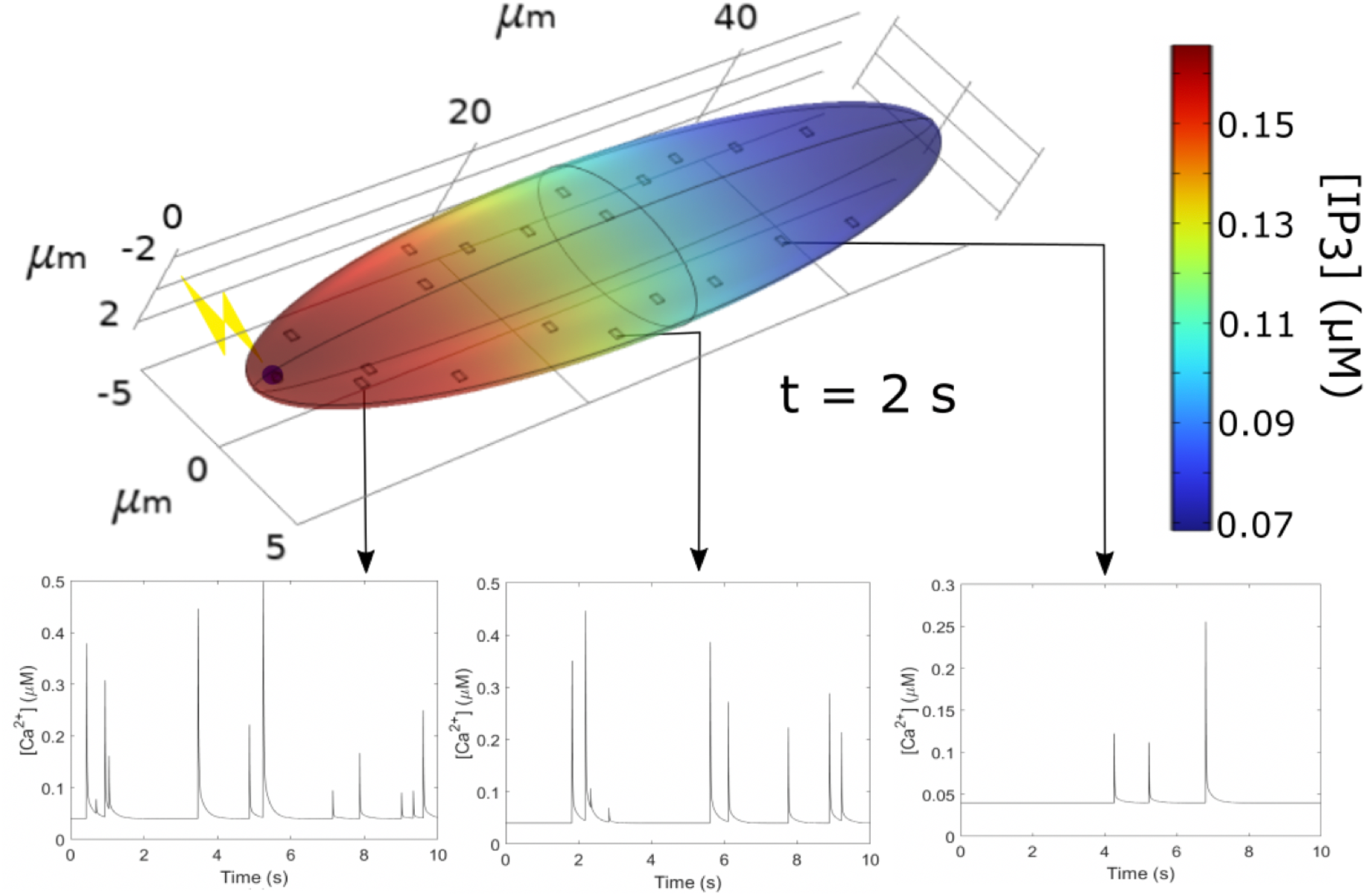
Computational simulations of IP_3_ diffusion and Ca^2+^ puff occurrence in response to the localized photorelease of a non-metabolizable IP_3_ analogue in an ellipsoidal 3D geometry, assuming an effective diffusion coefficient of IP_3_ (*D*_*I*_) of 100 µm^2^s^−1^. The upper panel shows IP_3_ distribution 2s after the flash. Lower panels show time series of local Ca^2+^ concentrations at cluster sites (in a 1 fL volume) located at increasing distances from the flash. Simulation procedures are the same as for Figure 4. The rate of localized IP_3_ photorelease, *θ*, is taken equal to 2500 µMs^−1^, which corresponds to the 250 µMs^−1^ value for the 2D case after volumetric adjustment.

The shape of the cell is expected to influence the rate of diffusion. We investigated this possible influence by simulating IP_3_ diffusion in a two-dimensional representation of an astrocyte (Figure 6). In response to a release of IP_3_ at the intersection between the cell body and one process, Ca^2+^ puffs occur sooner in the cell process than in the body. This is related to a faster IP_3_ diffusion in elongated structures where dilution is much reduced. For example, 2 s after the simulated flash, more elevated IP_3_ concentrations are seen in the process than in the cell body (Figure 6D). Thus, at the same time and the same distance from the site of IP_3_ release, IP_3_ reaches higher concentrations in the process and is able to propagate on longer distances (Supplemental Video S5).

**Figure 6.**
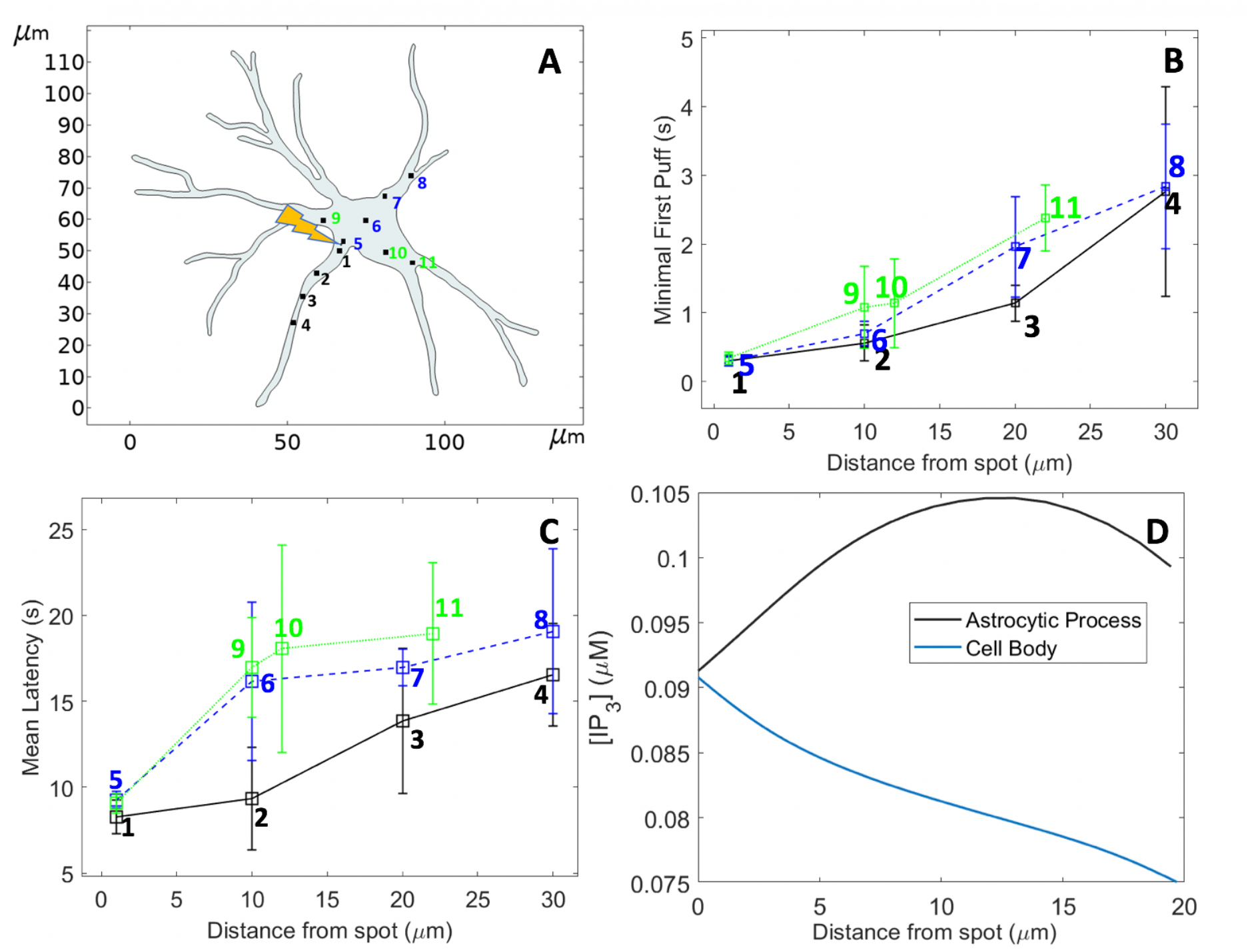
Computational simulations of IP_3_ diffusion and Ca^2+^ puff occurrence in response to the localized photorelease of a non-metabolizable IP_3_ analogue in a 2D geometry resembling an astrocyte, assuming an effective diffusion coefficient of IP_3_ (*D*_*I*_) of 100 µm^2^s^−1^. **A:** Shape of the astrocyte redrawn in COMSOL Multiphysics from images of Gonçalves-Pimentel et al. (2018). Locations of the clusters considered in the simulations are indicated and labelled in black in the astrocytic process and in blue and green in the cell body. IP_3_ photorelease was assumed to occur between locations 1 and 5. Simulation procedures are the same as for Figure 4 (main text), with *θ* = 250 µms^−1^ and flash duration = 500 ms. Panels **B and C** show the minimal first puffs and the mean first puff latencies at the different locations, respectively. Even at equal distance from the IP_3_ release point, puffs occur sooner in the process than in the body. Panel **D** shows the distribution of IP_3_ 2s after the flash: higher local IP_3_ concentrations are reached in the process in which there is no dilution effect. The spatio-temporal evolution of [IP_3_] can be seen in the Supplement Video S5.

In conclusion, the value inferred for the effective IP_3_ diffusion coefficient is not significantly affected by considering a 3D system instead of a 2D one. On the other hand, cell shape can considerably affect diffusion, with elongated geometries increasing concentration gradients and thereby favoring fast diffusion.

### Realistic ER geometry

The ER consists in a network of tubules and flattened sacs, resulting in a complex shape. Since the IP_3_ receptors are located within its membrane, diffusion of IP_3_ to the receptors may be affected by the ER structure. Taking advantage of the great flexibility of Comsol Multiphysics, we performed simulations in which the ER structure was explicitly considered. Thus, we simulated the geometry of a SH-SY5Y cell, in which we inserted an ER whose shape was largely inspired from 2D images of ER in DC-3F cells (De Angelis et al., 2019). We first compared the puff latencies simulated in the absence and in the presence of ER. The values of the rates of IP_3_ release were kept the same as in the simulations above (Figure 4), using a scaling factor to take into account the changes in the cytosolic volume. Puff latencies were in average not much affected by the presence of the ER (Figure 7). However, the time evolution of the corresponding IP_3_ profiles indicates that diffusion is locally affected by the extent and the shape of the accessible portions of cytosol between the IP_3_ release point and a given puff location. This is most easily visualized in an idealized, ellipse-shaped geometry. In this case, the IP_3_ profiles along a fictive line located at half the cell length and perpendicular to the gradient are indeed slightly different when IP_3_ is released at the right or the left extremity of the cell (Figure 8A-C). When compared to simulations where the ER structure is not considered, average IP_3_ increases are in average either a bit slower or similar (Figure 8D). However, locally, the presence of the ER can also allow for a faster IP_3_ increase (Figure 8B).

**Figure 7.**
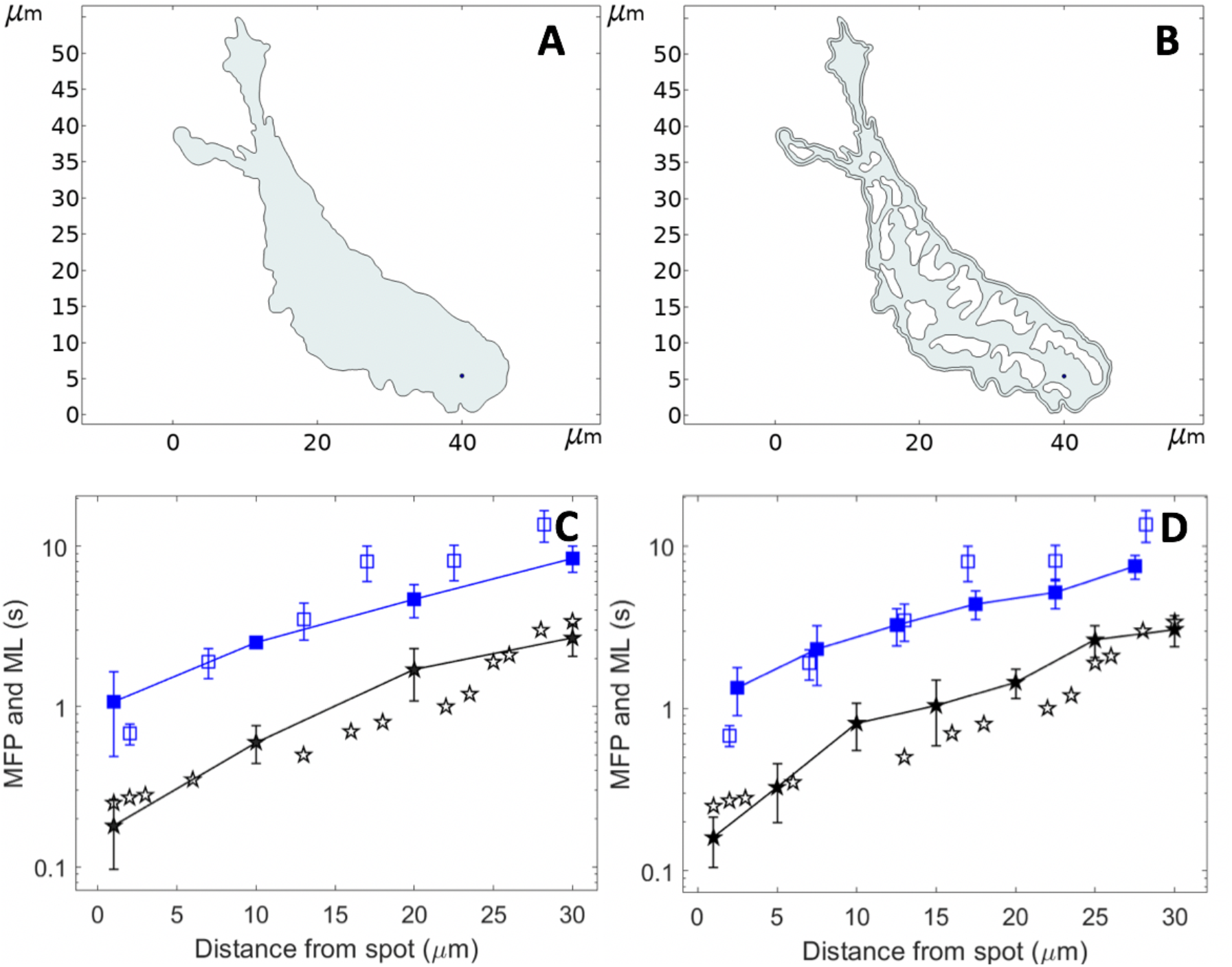
Theoretical investigation of the influence of the ER membranes on the diffusion of IP_3_ and Ca^2+^ puff occurrence. Panels **A and B** show the 2D geometry considered in the computational simulations of IP_3_ diffusion and Ca^2+^ puff occurrence in response to the localized photorelease of a non-metabolizable IP_3_ analogue. The shape of the cell was redrawn from Dickinson et al. (2016) and that of the ER was largely inspired from De Angelis et al. (2019). Panels **C and D** show the simulated (plain symbols) and experimental (empty symbols) mean latencies (blue) and minimal first puff latencies (black). Flash duration is 500 ms. Simulation procedures are the same as for Figure 4. The rates of localized IP_3_ photorelease, *θ*, is taken equal to 400.6 µMs^−1^ and 284.65 µMs^−1^, respectively which corresponds to the 250 µMs^−1^ used in the other simulations with the volumetric adjustments.

**Figure 8.**
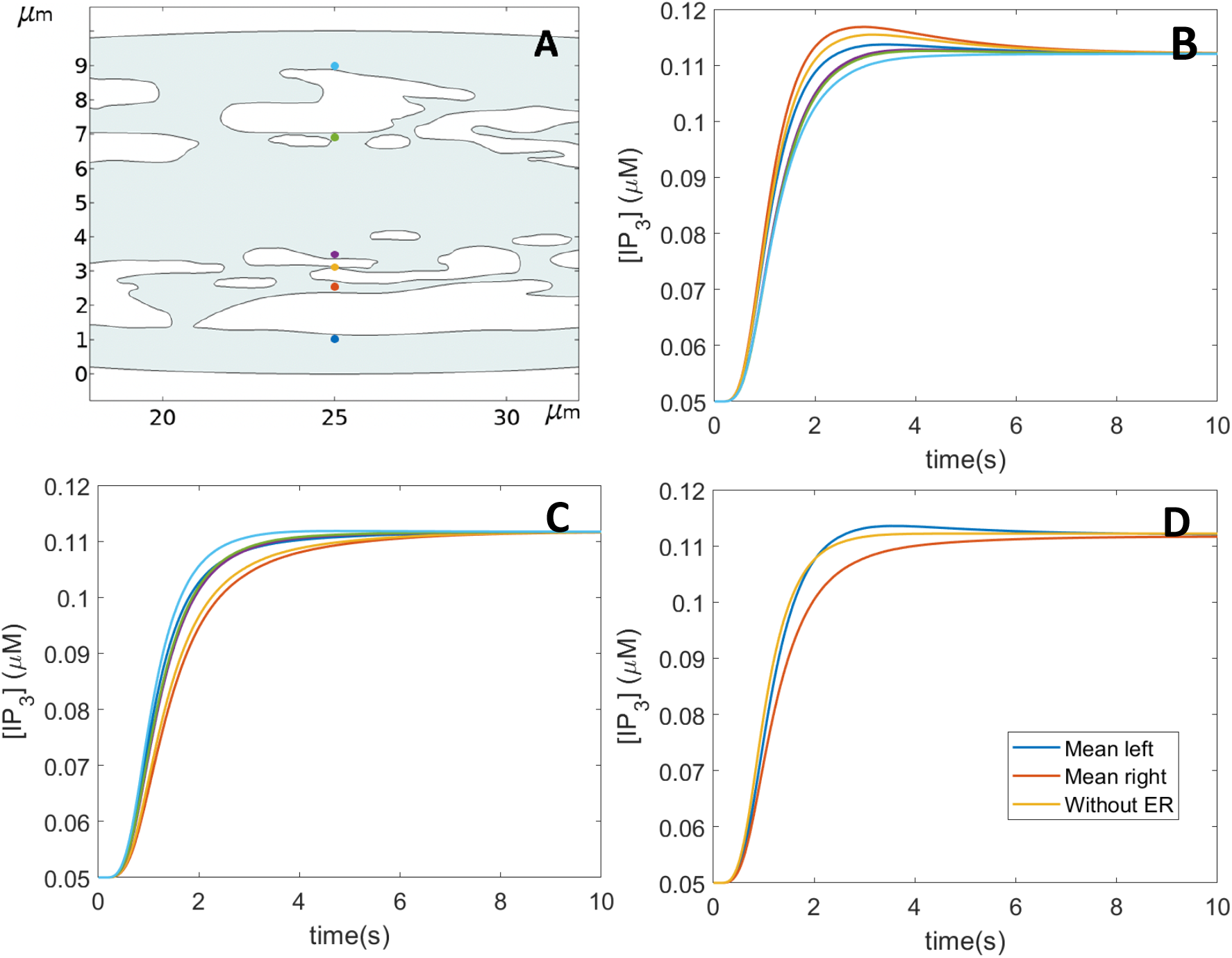
Theoretical investigation of the influence of the ER membranes on the diffusion of IP_3_ in a 2D ellipse-shaped cell (50 × 10 µm^2^). Panel **A** shows the central part of the simulated cell together with the locations of the clusters considered. Panels **B and C** show the IP_3_ temporal profiles at the different cluster locations considering a spot of photorelease of IP_3_ located at the left (B) or at the right (C) extremity of the cell. Line colors correspond to the points in panel A. In panel **D**, the profiles of the IP_3_ concentrations averaged on the six cluster sites are shown for the two situations corresponding to B and C. Simulation procedures are the same as for Figure 4. The rate of localized IP_3_ photorelease, *θ*, is taken equal to 179.42 µMs^−1^, which corresponds to the 250 µMs^−1^ used in the other simulations with the volumetric adjustments.

In conclusion, simulations indicate that the ER does not much affect average diffusion of IP_3_ in the cytosol. Although it constitutes a physical barrier that can locally slow down diffusion, it also creates cytosolic channel-like structures in which diffusion is accelerated because ions cannot spread in all directions.

## Discussion

IP_3_ plays a major role in Ca^2+^ signaling by mobilizing Ca^2+^ from the ER, which is the main intracellular Ca^2+^ store. After stimulation, IP_3_ must diffuse from the plasma membrane where it is produced across the cytoplasm to trigger Ca^2+^ release from the ER. Thus, IP_3_ diffusion plays an essential role in Ca^2+^ signaling. Following measurements in a medium devoid of IP_3_R, IP_3_ diffusion was assumed to be fast, with a diffusion coefficient of 283 ± 53 µm^2^s^−1^ (Allbritton et al., 1992). This estimation is in line with those of compounds of similar molecular weight such as ATP for example (Hubley et al., 1995). However, this value cannot account for the observation that in response to a localized release of a non-metabolizable analog of IP_3_, there is a ~10 s delay in the occurrence of Ca^2+^ puffs 30 nm away from the site of IP_3_ release (Dickinson et al., 2016). In the same manner, such a rapid diffusion cannot account for the drastic influence of the localization of the IP_3_-metabolizing 5-phosphatase enzyme on Ca^2+^ signaling observed in CHO cells by De Smedt et al. (1997). The latter authors developed a mutant InsP_3_ 5-phosphatase in which the C-terminal cysteine cannot be farnesylated, which hinders its binding to the plasma membrane. While the Ca^2+^ oscillations detected in the presence of 1 µM ATP were totally lost in 87.5% of intact (farnesylated) InsP_3_ 5-phosphatase-transfected cells, a loss of Ca^2+^ signal occurred in only 1.1% of the mutant InsP_3_ 5-phosphatase-transfected cells (De Smedt et al., 1997). Such a sensitivity to the location of an IP_3_ metabolizing enzyme could not be observed if IP_3_ was indeed a fast-diffusing molecule that rapidly becomes homogeneously distributed in the whole cell.

Accordingly, a much lower value of the effective diffusion coefficient of IP_3_ was predicted by Dickinson et al. (2016) on the basis of their observations of puff latencies in SH- SY5Y cells in response to the photorelease of a caged non metabolisable IP_3_ analog. On the basis of a simplified description of Ca^2+^ puff dynamics, these authors proposed that observations can be accounted for if the effective diffusion coefficient of IP_3_ in these cells is in the range 3-10 µm^2^s^−1^. This conclusion was re-examined in the present study, based on a detailed description of Ca^2+^ puff dynamics. Our results also point to a lower value of the effective diffusion coefficient of IP_3_ in intact cells than in cytosolic extracts of *Xenopus* oocytes but concluded to a decrease by a factor ~3 instead of 10 or more.

Besides the different computational framework, our approach differs from that developed by Dickinson et al. (2016) in three ways. First, only one spatial dimension was considered in the latter study, while we performed 2D or 3D simulations. As illustrated for simulations of IP_3_ diffusion in the 1D-like astrocytic process (Figure 6 and Video S5), diffusion is faster in 1D. The 1D approach of the previous study led to an underestimation of the diffusion coefficient because the value was fitted to reproduce the effective rate of IP_3_ diffusion that was observed in 3D. A second important difference relates to the relation between IP_3_ concentration and the probability of Ca^2+^ puff occurrence. At low IP_3_ concentration, the linear relation considered in the previous study predicts a larger probability of puff occurrence than the nonlinear function used in the present study (Equation 2). Again, this higher probability of puff occurrence was compensated by a lower value of the IP_3_ diffusion coefficient. Thirdly, because we here considered both the mean and the minimal first puff latencies, we inferred the rate of IP_3_ release by the flash (parameter θin Equations 3 and 5) in a more accurate way. We reasoned that the mean first puff latency may be affected by the global increase in basal Ca^2+^ if the latter is not fully prevented by the EGTA injected in the cell at a final concentration of 5 µM. In line with this hypothesis, agreement for both mean and first puff latencies was only obtained in our simulations when considering that the basal level of Ca^2+^ was in average 120 nM above basal level when puff activity was monitored after distributed photorelease of IP_3_. Because this Ca^2+^ increase was not taken into account in the previous study of Dickinson et al. (2016), the value of θwas overestimated, leading again to an underestimation of the IP_3_ diffusion coefficient.

To confirm that the IP_3_ diffusion coefficient is indeed larger than 10 µm^2^s^−1^, we simulated the experimental protocol of distributed photorelease of IP_3_ used in Dickinson et al. (2016) to induce a spatially uniform rise in IP_3_ concentration and described in the methodology section here above. Simulations performed with *D*_*I*_ = 10 µm^2^s^−1^ and shown in Figure S6B show that IP_3_ concentration remains spatially inhomogeneous up to at least 10 s after the flash. This is not in agreement with the observation that first puff latencies are independent from the location of the cluster under this protocol. In contrast, when *D*_*I*_ = 100 µm^2^s^−1^, IP_3_ rapidly equilibrates in the whole cell (Figure S6C).

Simulations predict that IP_3_ diffusion is not much affected by the proximity of the plasma membrane (compare results shown in Figures 4 and 5), nor by the precise shape of the cell. This is relevant because clusters of IP_3_ receptors are most of the time located very close to the plasma membrane (Smith et al., 2009b). However, elongated geometries favor fast diffusion and allow for higher local [IP_3_] than in extended systems. This was shown here by simulations in a situation corresponding to an astrocyte. Accordingly, rates of Ca^2+^ waves propagation in astrocytic processes are larger than in most cell types (33-100 µms^−1^, see Cornell-Bell and Finkbeiner, 1991 versus 10-20 µms^−1^, see Dupont et al., 2007). Surprisingly, the presence of the ER, which acts as an obstacle to free diffusion, does not much affect mean effective diffusion. This lack of global effect can be ascribed to the fact that while some paths are slower because of obstruction, others are faster because of the presence of channel-like structures which favor fast diffusion thanks to the absence of dilution effect. It should be noted that we did not considered the tubular-shaped of this network, nor its dynamical evolution into a more fragmented structure (Subramanian and Meyer, 1997).

The value of 100 µm^2^s^−1^ for the effective diffusion coefficient of IP_3_ that emerges from our simulations is in line with the buffering effect exerted by the non-fully bound IP_3_R tetramers, as proposed previously (Dickinson et al., 2016; Taylor and Konieczny, 2016). Thus, it is expected to be smaller in cell types with higher expression levels of IP_3_R. In the same line, effective diffusion is much accelerated upon increasing [IP_3_] because buffers become saturated. Thus, upon cell stimulation by an agonist that generally leads to a surge in [IP_3_] followed by a decrease, the properties of diffusion are expected to vary. Even more significant in this respect are the IP_3_ oscillations that have been observed in several cell types and that arise from the activation by Ca^2+^ of IP_3_ synthesis by PLC (Politi et al., 2006) and/or of IP_3_ metabolism by a 3- kinase (Dupont and Erneux, 1997; Sneyd et al., 2006). Further studies are required to assess the consequences of the interplay between the temporal changes in IP_3_ concentration and its diffusional properties.

## Acknowledgments

This work was supported by a PDR FRS-FNRS project (T.0073.21). GD is Research Director at the Belgian “Fonds National pour la Recherche Scientifique” (FRS-FNRS). We thank Benjamin Wacquier for scientific help and fruitful discussions, as well as Silvina Ponce Dawson for constructive comments.

## References

Allbritton N, Meyer T and Stryer L (1992) Range of messenger action of calcium ion and inositol 1,4,5-trisphosphate. Science 258: 1812–1815.

Alzayady K, Wang L, Chandrasekhar R, Wagner L, Van Petegem F and Yule D (2016) Defining the stoichiometry of inositol 1,4,5-trisphosphate binding required to initiate Ca2+ release. Science Signaling 9 (422): ra35.

Berridge MJ (1997) Elementary and global aspects of calcium signalling. Journal of Physiology 499: 291–306.

Bezprozvanny I, Watras J and Ehrlich B (1991) Bell-shaped calcium-response curves of Ins(1,4,5)P_3_- and calcium-gated channels from endoplasmic reticulum of cerebellum. Nature 351: 751–754.

Bootman M, Berridge M and Lipp P (1997) Cooking with calcium: The recipes for composing global signals from elementary events. Cell 91: 367–373.

Bootman M, Collins T, Peppiatt C, Prothero L, MacKenzie L, De Smet P, Travers M, Tovey S, Deo J, Berridge M, Ciccolini F and Lipp P (2001) Calcium signalling – an overview. Seminar Cell Developmental Biology 12: 3–10.

Calabrese A, Fraiman D, Zysman D and Ponce Dawson S (2010) Stochastic fire-diffuse-fire model with realistic cluster dynamics. Physical Review E 82: 031910.

Cornell-Bell A and Finkbeiner S (1991) Ca2+ waves in astrocytes. Cell Calcium 12: 185–204.

Dargan S, Parker I (2003) Buffer kinetics shape the spatiotemporal patterns of IP3-evoked Ca2+ signals. Journal of Physiology 553: 775–788.

De Angelis A, Denzi A, Merla C, Andre F, Garcia-Sanchez T, Mir L, Apollonio F and Liberti M (2019) Microdosimetric realisstic model of a cell with endoplasmic reticulum. Annual International Conference IEEE Engineering Medical Biological Society 2019: 134–137.

Decrock E, De Bock M, Wang N, Gadicherla A, Bol M, Delvaye T, Vandenabeele P, Vinken M, Bultynck G, Krysko D and Leybaert L (2013) IP_3_, a small molecule with a powerful message. Biochimica et Biophysica Acta 1833: 1772–1786.

De Smedt F, Missiaen L, Parys J, Vanweyenberg V, De Smedt H and Erneux C (1997) Isoprenylated human brain type I inositol 1,4,5-trisphosphate 5-phosphatase controls Ca2+ oscillations induced by ATP in Chinese hamster ovary cells. Journal of Biological Chemistry 272: 17367–17375.

Dickinson G, Swaminathan D, Parker I (2012) The probability of triggering calcium puffs is linearly related to the number of inositol trisphosphate receptors in a cluster. Biophysical Journal 102: 1826–1836.

Dickinson G, Ellefsen K, Ponce Dawson S, Pearson J and Parker I (2016) Hindered cytoplasmic diffusion of inositol trisphosphate restricts its cellular range of action. Science Signaling 9: ra108.

Dupont G and Erneux C (1997) Simulations of the effects of inositol 1,4,5-trisphosphate 3-kinase and 5-phosphatase activities on Ca2+ oscillations. Cell Calcium 22: 321–331.

Dupont G and Dumollard R (2004) Simulation of calcium waves in ascidian eggs: insights into the origin of the pacemaker sites and the possible nature of the sperm factor. Journal of Cell Science 117: 4313–4323.

Dupont G, Combettes L and Leybaert L (2007) Calcium dynamics: spatio-temporal organization from the subcellular to the organ level. International Review in Cytology 261: 193–245.

Dupont G, Falcke M, Kirk V and Sneyd J (2016) Models of calcium signalling. Interdisciplinary Applied Mathematics (Springer International Publishing, Switzerland, 2016), Vol 43.

Dupont G, Abou-Lovergne A and Combettes L (2008) Stochastic aspects of oscillatory Ca2+ dynamics in hepatocytes. Biophysical Journal 95: 2193–2202.

Gonçalves-Pimentel C, Moreno GMM, Trindade BS, Isaac AR, Rodrigues CG, Savarira M, de Albuquerque AV, de Andrade Aguiar JL, Andrade-da-Costa Blds (2018) Cellulose exopolysaccharide from sugarcane molasses as a suitable substrate for 2D and 3D neuron and astrocyte primary cultures. Journal of Material Science: Materials in Medecine 29: 139.

Hubley M, Moerland T and Rosanke R (1995) Diffusion coefficient of ATP and creatine phosphate in isolated muscle: pulsed gradient 31P NMR of small biological sample. NMR in Biomedecine 8: 72–78.

Keebler M and Taylor C (2017) Endogenous signalling pathways and caged IP3 evoke Ca2+ puffs at the same abundant immobile intracellular sites. Journal of Cell Science 130: 3728–3739.

Leybaert L (2016) IP3, still on the move but now in the slow lane. Science Signaling 9: fs17.

Lock J and Parker I (2020) IP3 mediated global Ca2+ signals arise through two temporally and spatially distinct modes of Ca2+ release. eLife 9: e55008.

Luzzi V, Sims C, Soughayer J and Allbritton N (1998) The physiological concentration of inositol 1,4,5-trisphosphate in the oocytes of Xenopus laevis. Journal of Biological Chemistry 273: 28657–28662.

Politi A, Gaspers L, Thomas A and Höfer T (2006) Models of IP3 and Ca2+ oscillations: frequency encoding and identification of underlying feedbacks. Biophysical Journal 90: 3120–3133.

Sammels E, Parys J, Missiaen L, De Smedt H and Bultynck G (2010) Intracellular Ca2+ storage in health and disease: a dynamic equilibrium. Cell Calcium 47: 297–314.

Skupin A, Kettenman H, Winkler U, Wartenberg M, Sauer H, Tovey S, Taylor C and Falcke M (2008) How does intracellular Ca2+ oscillate: By chance or by the clock? Biophysical Journal 94: 2404–2411.

Smith I and Parker I (2009a) Imaging the quantal substructure of single IP3R channel activity during Ca2+ puffs in intact mammalian cells. Proceedings of the National Academy of Science USA 106: 6404–6409.

Smith I, Wiltgen S and Parker I (2009b). Localization of puff sites adjacent to the plasma membrane: functional and spatial characterization of Ca2+ signaling in SH-SY5Y cells utilizing membrane-permeant caged IP3. Cell Calcium 45: 65–76.

Sneyd J, Tsaneva-Atanasova K, Reznikov V, Bai Y, Sanderson M and Yule D (2006) A method for determining the dependence of calcium oscillations on inositol trisphosphate oscillations. Proceedings of the National Acadamy of Sciences USA 103: 1675–1680.

Spät A, Bradford P, McKinney J, Rubin R and Putney J (1986) A saturable receptor for 32P-inositol-1,4,5-trisphosphate in hepatocytes and neutrophils. Nature 319: 514–516.

Subramanian K and Meyer T (1997) Calcium-induced restructuring of nuclear envelope and endoplasmic reticulum calcium stores. Cell 89: 963–971.

Swillens S, Dupont G, Combettes L and Champeil P (1999) From calcium blips to calcium puffs: theoretical analysis of the requirements for interchannel communication. Proceedings of the National Academy of Science USA 96: 13750–13755.

Tanimura A, Morita T, Nezu A, Shitara A, Hashimoto N and Tojyo Y (2009) Use of fluorescence resonance energy tansfer-based biosensors for the quantitative analysis of inositol 1,4,5-trisphosphate dynamics in calcium oscillations. Journal of Biological Chemistry 284: 8910–8917.

Taylor CW, Konieczny V (2016) IP3 receptors: Take four IP3 to open. Science Signaling 9(422):pe1.

Thurley K, Tovey S, Moenke G, Prince V, Meena A, Thomas A, Skupin A, Taylor C and Falcke M (2014) Reliable encoding of stimulus intensities with random sequences of intracellular Ca2+ spikes. Science Signaling 7: ra59.

Tovey S, de Smet P, Lipp P, Thomas D, Young K, Missiaen L, De Smedt H, Parys J, Berridge M, Thuring J, Holmes A and Bootman D (2001) Calcium puffs are generic InsP3-activated elementary calcium signals and are downregulated by prolonged hormonal stimulation to inhibit cellular calcium responses. Journal of Cell Science 114: 3979–3989.

Thurley K, Tovey S, Moenke G, Prince V, Meena A, Thomas A, Skupin A, Taylor C and Falcke M (2014) Reliable encoding of stimulus intensities within random sequences of intracellular Ca2+ spikes. Science Signaling 7(331): ra59.

Voorsluijs V, Ponce Dawson S, De Decker Y and Dupont G (2019) Deterministic limit of intracellular calcium spikes. Physical Review Letters 122: 088101.

Wojcikiewicz R (1995) Type I, II, and III inositol 1,4,5-trisphosphate receptors are unequally susceptible to down-regulation and are expressed in markedly different proportions in different cell types. Journal of Biological Chemistry 270: 11678–11683.

Yao Y, Choi, I, Parker, I (1995) Quantal puffs of intracellular Ca2+ evoked by inositol trisphosphate in Xenopus oocytes. Journal of Physiology 482: 533–553.

