## Supplementary material for "Computational investigation of IP_3_ diffusion": Supp

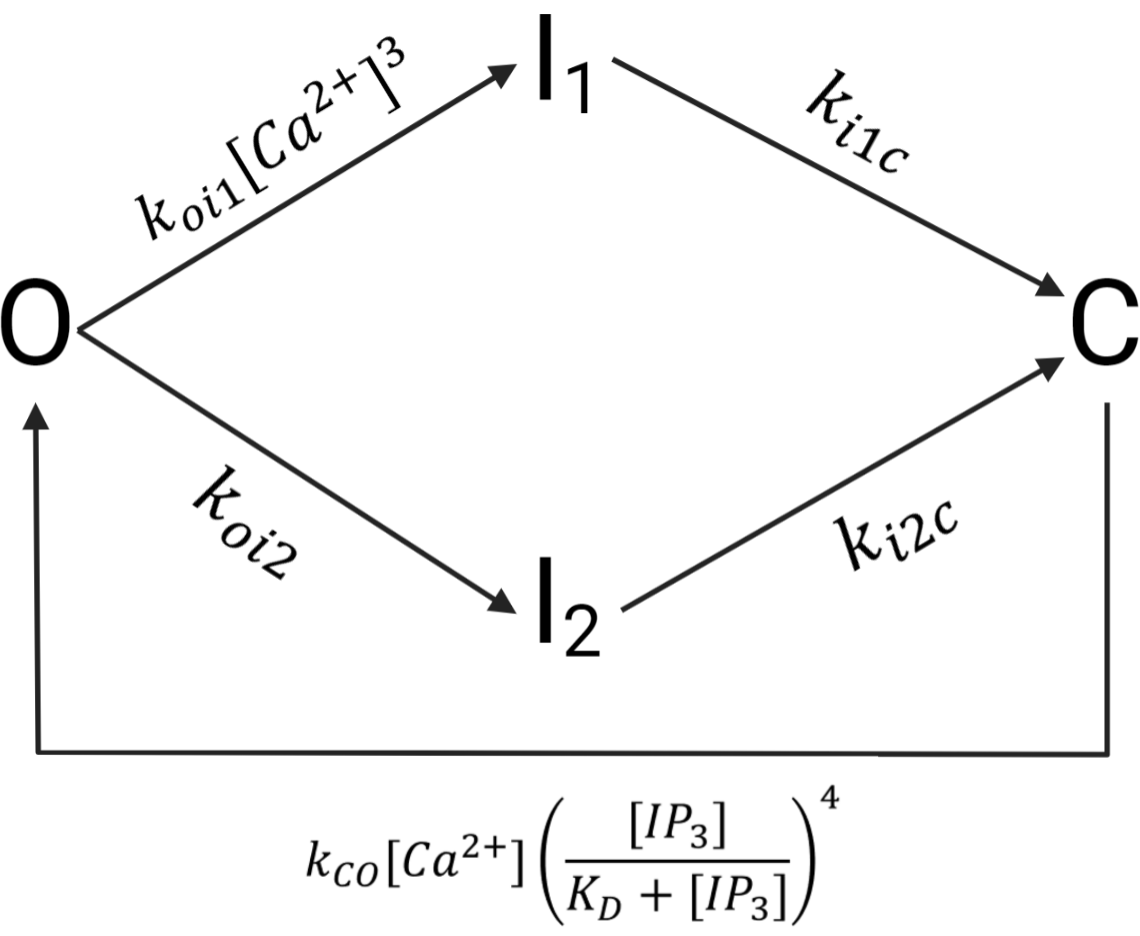

Figure S1

Schematic representation of the model used to simulate the dynamics of a cluster of IP_3_R’s. The model is taken from Calabrese et al. (2010), where it was calibrated against experimental observations. Here, we explicitly incorporate the dependence of the rate of passage of the cluster from the closed (C) to the open (O) state on [IP_3_].

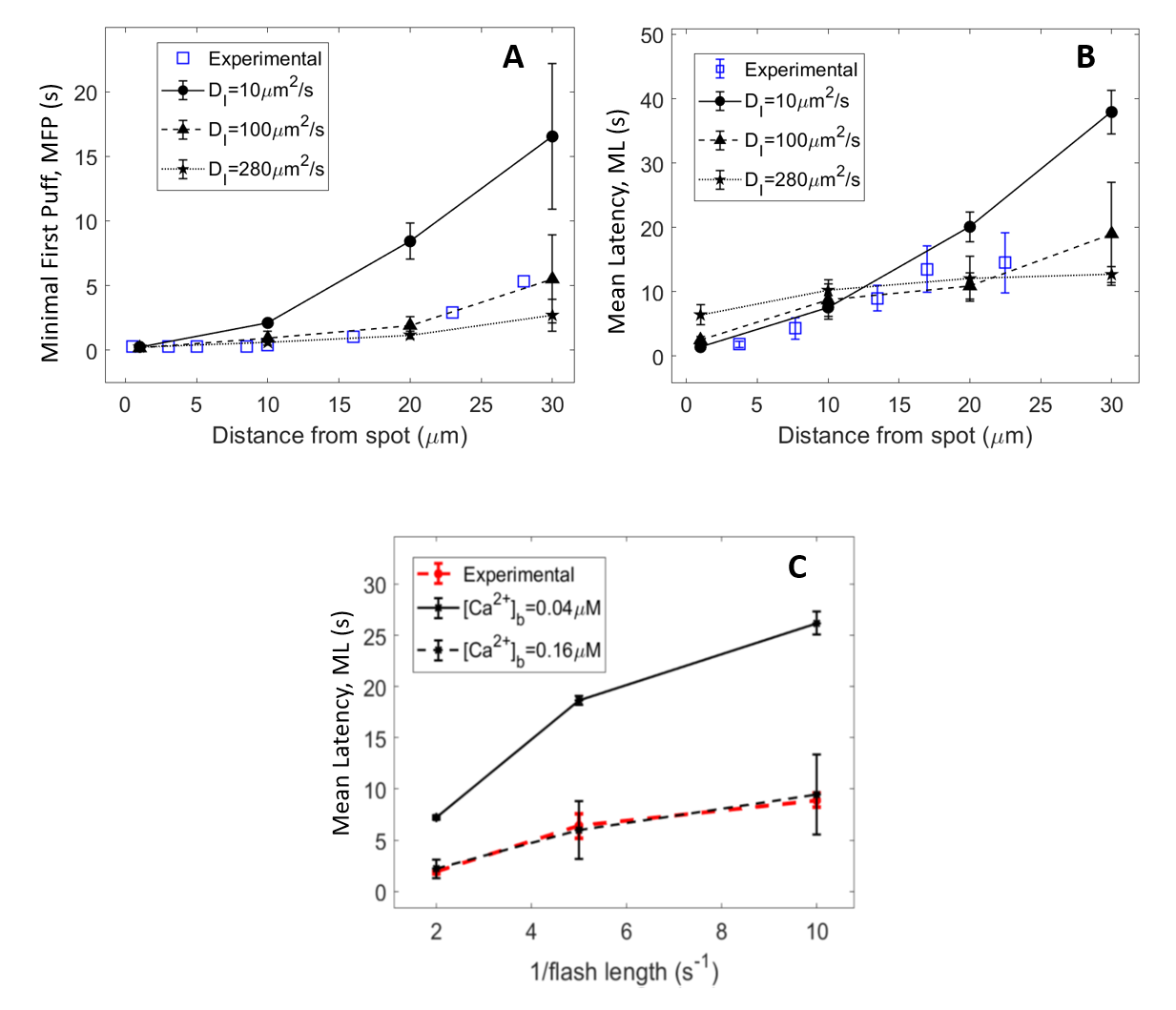

Figure S2.

Comparison between puff latencies theoretically predicted with the rate of IP_3_ release *θ* = 250 μm^2^s^-1^ and observed by Dickinson et al. (2016). Panels **A and B** show minimal first puff latencies and mean latencies simulated with different values of the effective diffusion coefficient for IP_3_, *D_I_*. All points correspond to a 200 ms flash. Plain symbols are theoretical predictions while empty symbols are the experimental observations of Dickinson et al. (2016). Lines are drawn between simulation results. Simulations are performed in a 2D ellipse-shaped system with the spot of IP_3_ release being a 0.25 μm radius disk at one extremity of the simulated cell. For each theoretical point, 50 independent simulations were run. Error bars indicate ± SEM. For minimal first puffs, the 50 simulations were divided in 10 groups of 5 simulations among which the minimal first puff was considered.

Panel **C** shows the mean latency as a function of the inverse of flash duration for the re-estimated value of *θ* (600 μMs^-1^) for two different values of basal Ca^2+^ concentration. The increase in basal Ca^2+^ concentration is thought to arise from the uncomplete Ca^2+^ buffering by EGTA in the experiments and explains why the value of *θ* directly inferred from the experiments with distributed photorelease of IP_3_ is overestimated.

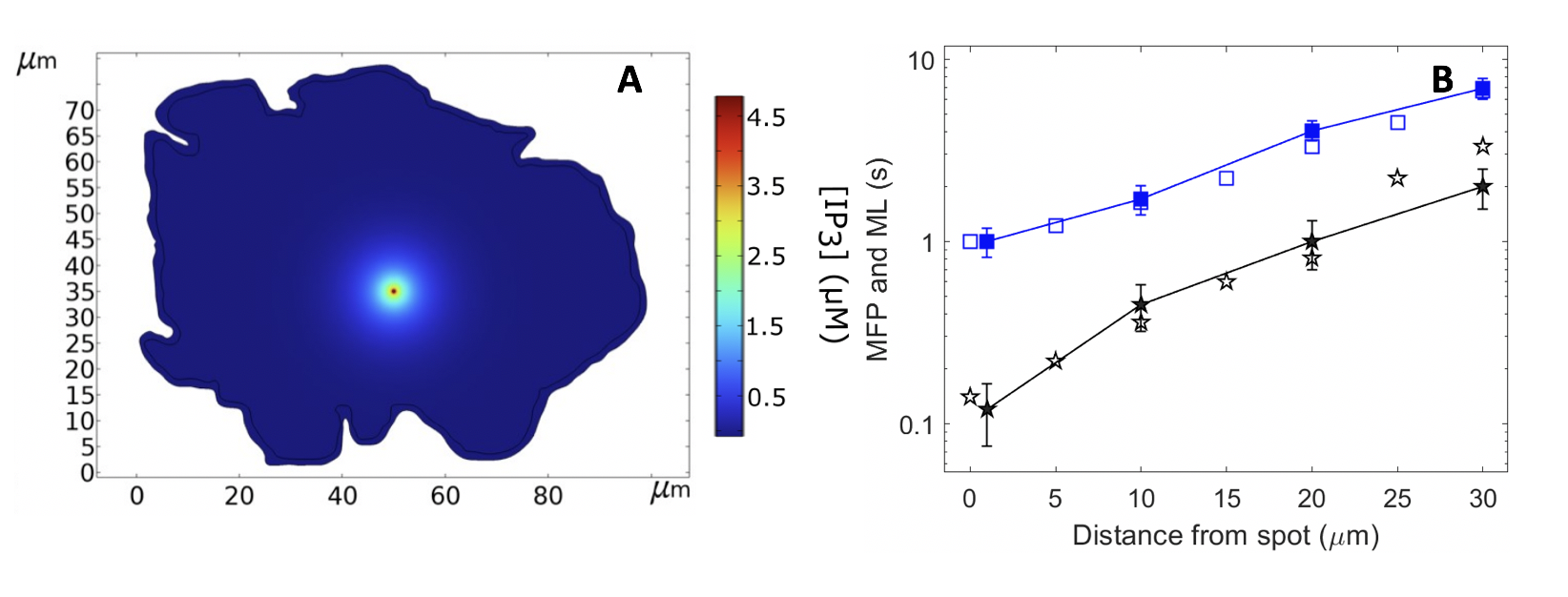

Figure S3

Simulations of Ca^2+^ puff occurrence in response to the localized photorelease of a non-metabolizable IP_3_ analog in COS-7 cells, assuming an effective diffusion coefficient of IP_3_ *D_I_* = 100 μm^2^s^-1^. Panel **A** shows the simulated cell, redrawn in COMSOL Multiphysics, with the distribution of IP_3_ concentration at the end the flash of the IP_3_ analog (θ = 3591.73 μMs^-1^). Panel **B** shows simulated (plain dots) and experimental (empty dots) mean latencies (blue) and minimal first puff latencies (black). Flash duration is 500 ms. The shape of the cell and the experimental values of latencies are taken from Dickinson et al. (2016). Simulation procedures are the same as for Figure 3 (main text).

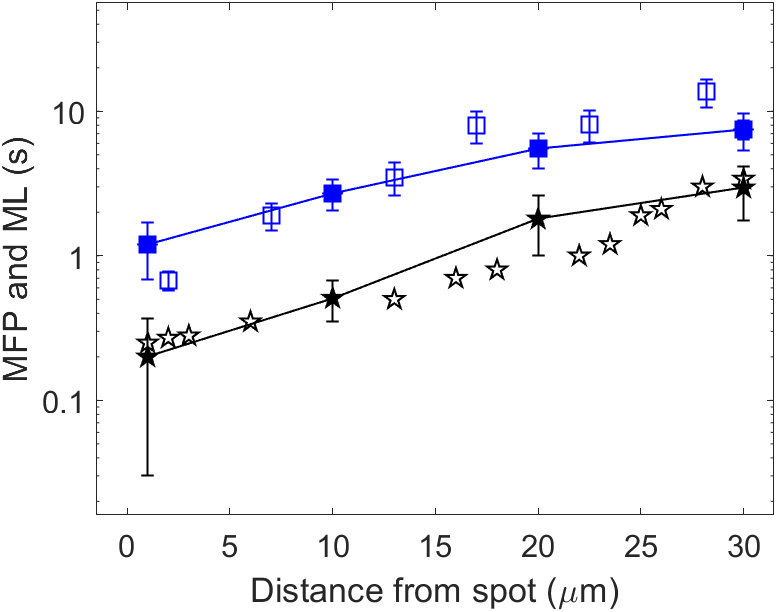

Figure S4

Simulations of Ca^2+^ puff occurrence in response to the localized photorelease of a non-metabolizable IP_3_ analog in an ellipsoidal 3D geometry, assuming an effective diffusion coefficient of IP_3_ *D_I_* = 100 μm^2^s^-1^. Results correspond to the spatio-temporal simulations shown in Figure 4. Shown are the simulated (plain symbols) and experimental (empty symbols) mean latencies (blue) and minimal first puff latencies (black). Flash duration is 500 ms. Experimental values of latencies are taken from Dickinson et al. (2016). Simulation procedures are the same as for Figure 3 (main text). The rate of localized IP_3_ photorelease, *θ* , was taken equal to 2500 μMs^-1^, which corresponds to the 250 μMs^-1^ value for the 2D case.

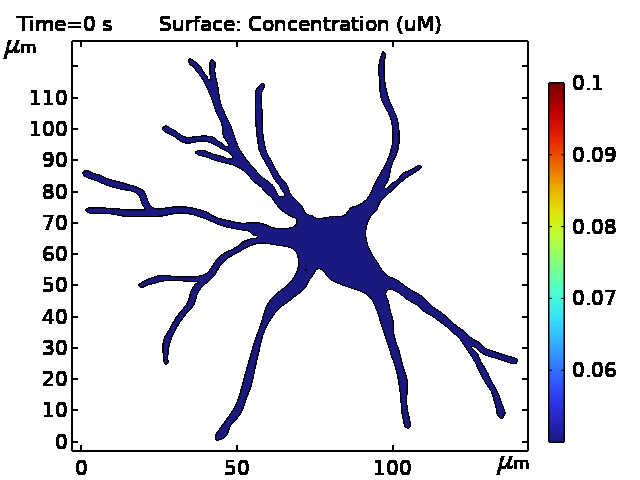

Video S5

Simulations of IP_3_ diffusion in response to the localized photorelease of a non-metabolizable IP_3_ analogue in a 2D geometry resembling an astrocyte, assuming an effective diffusion coefficient of IP_3_ *D_I_* = 100 μm^2^s^-1^. Simulation procedure is described in the legend of Figure 5 (main text).

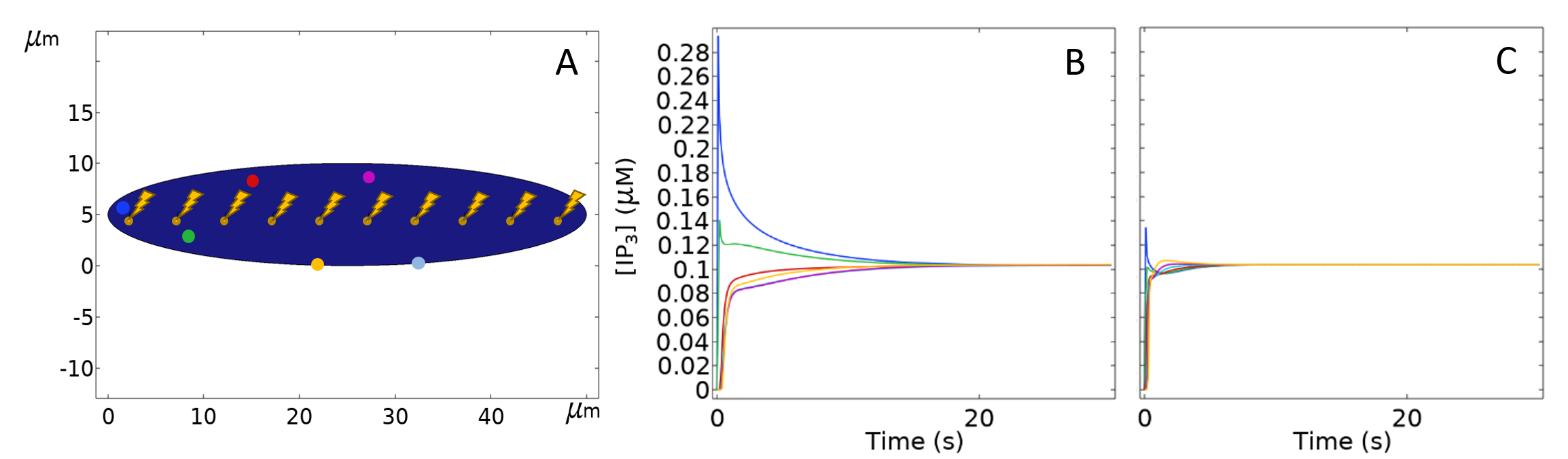

Figure S6

Simulations of the protocol of distributed photorelease used by Dickinson et al. (2016). As illustrated in panel **A**, IP_3_ is liberated at 10 different spots to provoke an increase that is supposed to be nearly homogenous in the whole cell, which is indeed indicated by the observation that puff latencies do not depend on the spatial location of the puff sites in these conditions. To accurately reproduce the experimental protocol, IP_3_ is sequentially released during 50 ms at a rate *θ* = 250 μMs^-1^, at each spot from left to right. The total amount thus corresponds to a 500 ms flash of 25 μMs^-1^ intensity, as modelled in Figure 4. Panel **B** shows the time evolutions of the IP_3_ concentrations at the locations of the dots of corresponding color in panel A, when *D_I_* = 10 μm^2^s^-1^. Panel **C** shows the time evolutions of the IP_3_ concentrations at the locations of the dots of corresponding color in panel A, when *D_I_* = 100 μm^2^s^-1^.

Table S1

Propensities and evolution equations used in the simulations of puffs and spikes dynamics.

| **Event** | **Propensity functions** $\mathbf{(}\boldsymbol{\omega}_{\boldsymbol{r}}\mathbf{)}$ | **Action** |
| --- | --- | --- |
| $\boldsymbol{C}\boldsymbol{\to}\boldsymbol{O}$ | $k_{co}\frac{N_{ca}}{\Omega}\left( \frac{[{IP}_{3}]}{K_{D}+\left[ {IP}_{3} \right]} \right)^{4}\left( 1-N_{o}-N_{i1}-N_{i2} \right)$ | $N_{c}=0$  $N_{o}=1$ |
| $\boldsymbol{O}\boldsymbol{\to}\boldsymbol{I}_{\mathbf{1}}$ | $k_{oi1}N_{o}\frac{N_{ca}\left( N_{ca}-1 \right)\left( N_{ca}-2 \right)}{\Omega^{3}}$ | $N_{o}=0$  $N_{i1}=1$ |
| $\boldsymbol{O}\boldsymbol{\to}\boldsymbol{I}_{\mathbf{2}}$ | $k_{oi1}N_{o}$ | $N_{o}=0$  $N_{i2}=1$ |
| $\boldsymbol{I}_{\mathbf{1}}\boldsymbol{\to}\boldsymbol{C}$ | $k_{i1c}N_{i1}$ | $N_{i1}=0$  $N_{c}=1$ |
| $\boldsymbol{I}_{\mathbf{2}}\boldsymbol{\to}\boldsymbol{C}$ | $k_{i2c}N_{I2}$ | $N_{i2}=0$  $N_{c}=1$ |

**Equations describing Ca^2+^ diffusion and Ca^2+^ handling mechanisms**

At a cluster site

In the cytoplasm

$$\frac{\partial\left[ {Ca}^{2+} \right]}{\partial t}=D_{C}\nabla^{2}\left[ {Ca}^{2+} \right]+J_{leak}-J_{SERCA}$$

$$\frac{\partial\left[ {Ca}^{2+} \right]}{\partial t}={D_{C}\nabla}^{2}\left[ {Ca}^{2+} \right]+\text{Σ}\text{o}$$

$$J_{SERCA}=\frac{v_{p}\left[ {Ca}^{2+} \right]^{2}}{{K_{p}}^{2}+\left[ {Ca}^{2+} \right]^{2}}$$

$$J_{leak}=\frac{v_{p}\left[ {Ca}^{2+} \right]_{b}^{2}}{{K_{p}}^{2}+\left[ {Ca}^{2+} \right]_{b}^{2}}$$

Table S2

Default values of the parameters used in all simulations except when mentioned explicitly.

| Parameter | Definition | Value |
| --- | --- | --- |
| k_co_ | C→O | 50 µM^-1^s^-1^ |
| k_oi1_ | O→I_1_ | 0.05 µM^-3^s^-1^ |
| k_oi2_ | O→I_2_ | 40 s^-1^ |
| k_i1c_ | I_1_→C | 0.005 s^-1^ |
| k_i2c_ | I_2_→C | 2 s^-1^ |
| ν_p_ | Maximal rate of SERCA | 0.9 µM s^-1^ |
| K_p_ | SERCA binding constant | 0.1 µM |
| [Ca^2+^]_b_ | Basal [Ca^2+^] | 0.04 µM |
| [IP_3_]_b_ | Basal [IP_3_] | 0.05 µM |
| D_C_ | Ca^2+^ diffusion coefficient | 40 µm^2^s^-1^ |
| Σ | Ca^2+^ release rate from one cluster | 500 µM s^-1^ |
| K_D_ | IP_3_ dissociation constant from IP_3_Rs | 0.1µM |
| L_c_ | Length of a compartment in the simulations | 0.5 μm |
| V_c_ | Volume of a compartment in the simulations | 10^-16^ L |
| Ω | Extensivity parameter | N_AV_ . V_c_. 10^-6^ |
